# CHIME: CMOS-hosted in-vivo microelectrodes for massively scalable neuronal recordings

**DOI:** 10.1101/570069

**Authors:** Mihaly Kollo, Romeo R Racz, Mina-Elraheb S Hanna, Abdulmalik M Obaid, Matthew R Angle, William Wray, Yifan Kong, Andreas Hierlemann, Jan Müller, Nicholas A Melosh, Andreas T Schaefer

**Affiliations:** Neurophysiology of Behaviour Laboratory, Francis Crick Institute, NW1 1AT, London, UK; Department of Neuroscience, Physiology & Pharmacology, University College London; Department of Materials Science and Engineering, Stanford University, Stanford, CA, USA; ETH Zurich, Department Biosystems Science and Engineering, Basel, Switzerland; Paradromics Inc., Austin, TX, USA; MaxWell Biosystems AG, Mattenstrasse 26, Basel, Switzerland

## Abstract

Mammalian brains consist of 10s of millions to 100s of billions of neurons operating at millisecond time scales, of which current recording techniques only capture a tiny fraction. Recording techniques capable of sampling neural activity at such temporal resolution have been difficult to scale: The most intensively studied mammalian neuronal networks, such as the neocortex, show layered architecture, where the optimal recording technology samples densely over large areas. However, the need for application-specific designs as well as the mismatch between the threedimensional architecture of the brain and largely two-dimensional microfabrication techniques profoundly limits both neurophysiological research and neural prosthetics.

Here, we propose a novel strategy for scalable neuronal recording by combining bundles of glass-ensheathed microwires with large-scale amplifier arrays derived from commercial CMOS of in-vitro MEA systems or high-speed infrared cameras. High signal-to-noise ratio (<20 μV RMS noise floor, SNR up to 25) is achieved due to the high conductivity of core metals in glass-ensheathed microwires allowing for ultrathin metal cores (down to <1 μm) and negligible stray capacitance. Multi-step electrochemical modification of the tip enables ultra-low access impedance with minimal geometric area and largely independent of core diameter. We show that microwire size can be reduced to virtually eliminate damage to the blood-brain-barrier upon insertion and demonstrate that microwire arrays can stably record single unit activity.

Combining microwire bundles and CMOS arrays allows for a highly scalable neuronal recording approach, linking the progress of electrical neuronal recording to the rapid scaling of silicon microfabrication. The modular design of the system allows for custom arrangement of recording sites. Our approach of employing bundles of minimally invasive, highly insulated and functionalized microwires to lift a 2-dimensional CMOS architecture into the 3rd dimension can be translated to other CMOS arrays such as electrical stimulation devices.

## Introduction

There is an increasing interest in understanding large nervous systems, such as the brains of mammals including humans. For these investigations, the main obstacle is the large size of the brain, and the extensive population coding employed, that makes it essential to monitor the activity of large distributed neuronal ensembles in most behavioural states (Georgopoulos et al., 1986; Lee et al., 1988; Sanger, 2003; Averbeck et al., 2006; Miura et al., 2012; Gschwend et al., 2016). Brain-machine interfaces (BMI) in turn seek to query and control the state of neuronal networks and have great promise for neuronal prosthetics. It is clear that for efficient measurement and control, the number of monitored units needs to extend way beyond the currently available methods (Chapin, 2004).

Electrical recordings are the state-of the art for BMI and prosthetics, can record at millisecond timescales, and have been used to achieve the largest channel-count recordings capable of time-resolving action potentials (Buzsáki, 2004; Du et al., 2011; Berényi et al., 2014; Shobe et al., 2015; Jun et al., 2017). Despite these advances, the silicon-based approaches for large-scale recordings have several limitations: Silicon probes cover only limited space, and recording sites are generally arranged inflexibly along a vertical/axial direction, while in the most studied brain areas (cortex, hippocampus, olfactory bulb), sampling in broad horizontal layers is best for studying representations. Device integration challenges have so far prevented their insertion at high lateral density <400 μm inter-probe distance. Another limiting factor is that incremental iterations in the design are typically subject to long turnaround times associated with ASIC verification, validation, and fabrication. Finally, while each electrode is small, the size of the overall shank and resulting tissue damage is posing a great limit to scaling by using multiple silicone probes (Buzsáki, 2004; Kozai et al., 2012; Woolley et al., 2013; Prodanov and Delbeke, 2016).

Designing a massively scalable electrophysiological recording system poses challenges at multiple levels and requires a systematic design effort (Figure 1). Apart from the individual electrodes, scaling has been frustrated by the engineering challenge for connectors, amplification and digitization. One approach to mitigate this has been to integrate electronics, multiplexing before connecting (Lopez et al., 2014; Raducanu et al., 2017; Fiáth et al., 2018), but integrating active electronics also requires a largely planar electrode architecture. A further limit to upscaling these approaches is that with further size reduction of individual electrodes, generally coupling impedance and signal loss increase. A potential solution that was suggested is to incorporate active electronics at the recording site (Neves et al., 2008; Lopez et al., 2014; Ruther and Paul, 2015). However, space constraints and temperature limit the scalability with this approach (Marblestone et al., 2013).

**Figure 1.**
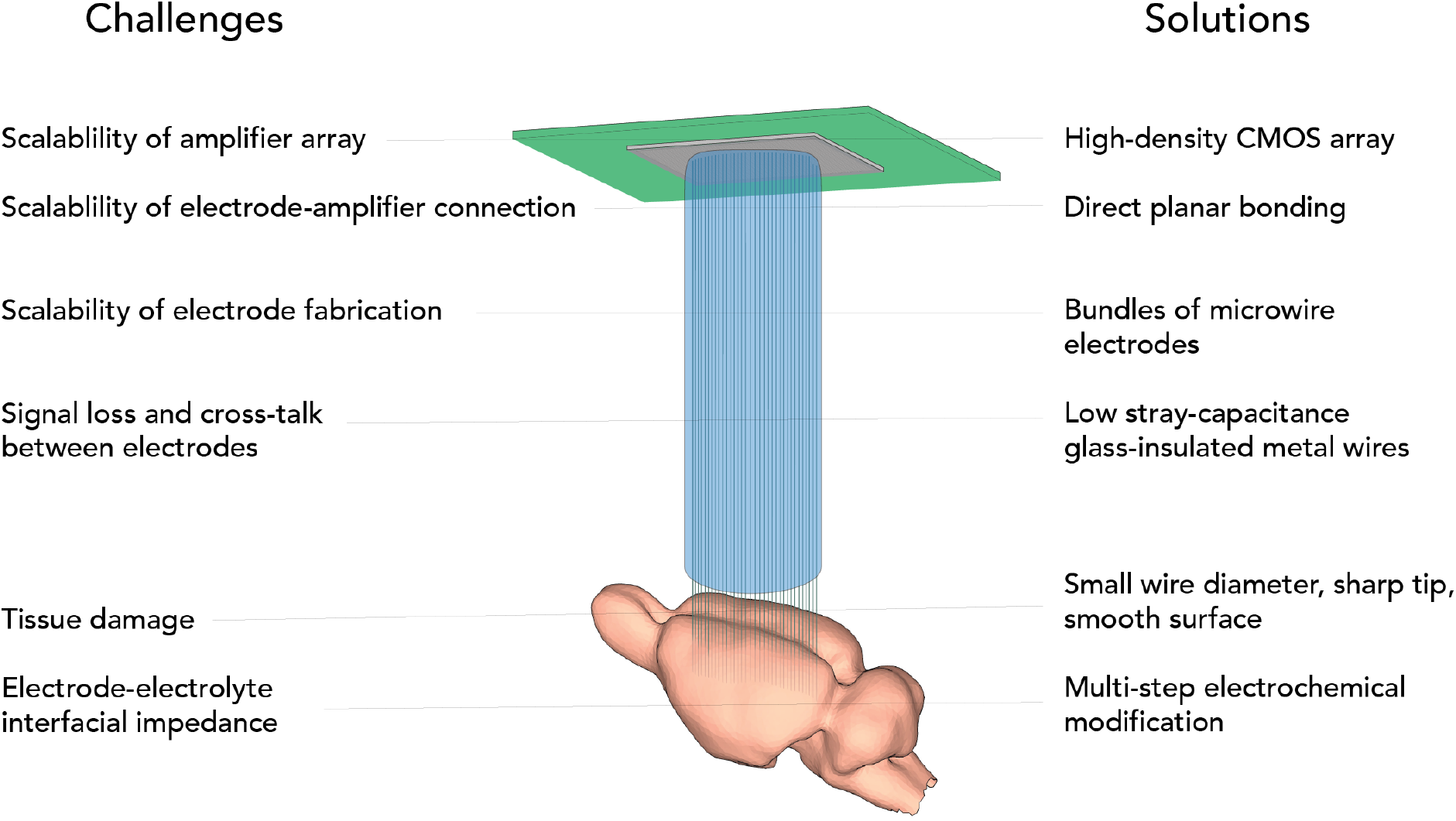
Challenges of scalable electrophysiology addressed by CHIME.

All these factors together might underlie the unfortunate situation that scaling has been extremely slow compared to other technology (Stevenson and Kording, 2011) and this speaks to the need for fundamentally new approaches (Marblestone et al., 2013). For slice and culture recordings on the other hand, scaling has advanced more rapidly, by making use of CMOS-based approaches (Scribner et al., 2007; Ferrea et al., 2012; Müller et al., 2015; Dragas et al., 2017; Tsai et al., 2017), however these techniques are inherently only suitable for flat samples, and not for in-vivo brain recordings in large brains, such as rodents or primates.

Here we propose to make use of the favourable scaling properties of CMOS devices and lift the flat CMOS architecture into the third dimension using ultra low impedance microwire bundles, thus making it suitable for brain recordings (Figure 1). We term this approach CMOS-Hosted In vivo MicroElectrode (CHIME) recordings.

## Results

### Microwire bundle fabrication

An ideal electrode for scalable neuronal recordings has good signal-to-noise characteristics, penetrates into the tissue without causing significant damage, while increased number of channels can be achieved without excessive fabrication effort. Volume displacement, and friction can impact on the tissue-electrode interaction upon insertion (Kozai et al., 2010, 2012). Therefore, we postulate that ideal microwires should be small, sharp and smooth. At the same time, to achieve large signal-to-noise ratios, recordings should be focal, where the geometric electrode area is below a few μm^2^ (Teleńczuk et al., 2018). To achieve all this, we used a modified Taylor-Ulitovsky method (Zhukov, 2006; Baranov et al., 2017) that allows fabrication of kilometre-long, glass insulated microwires with conductive metal cores (Figure 2A).

**Figure 2.**
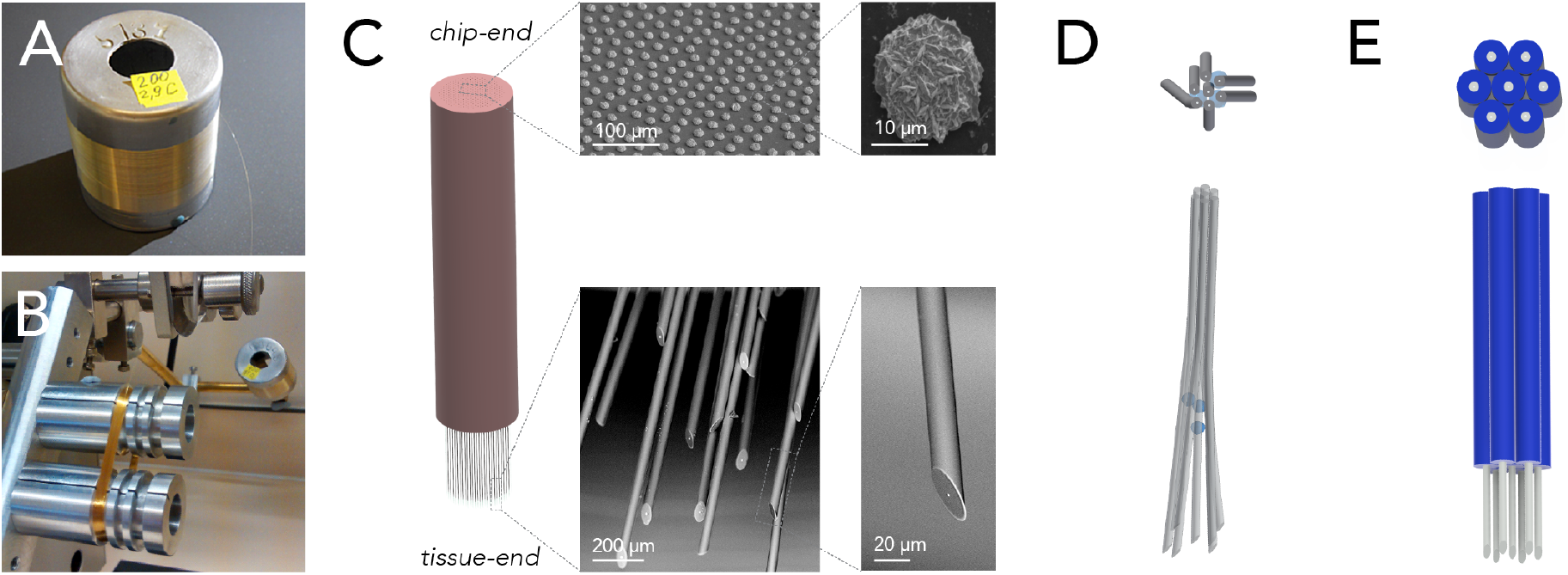
Scalable fabrication of CHIME electrode bundles. (A) Cast glass-insulated metal microwires are produced in several km long, continuously conductive stretches. (B) The flexibility and robustness of the composite wires allow the preparation of large wire-count bundles by spooling. The image shows a 1000 wire bundle prepared on a fibre winding machine. (C) The bundle is embedded into polymer, and both ends of the fibre bundle are altered with scalable procedures. For optimal contact with the electronics, the chip-end of the embedded bundle is polished flat and the exposed metal conductor cores at the chip-end of the wires are increased by electrochemical gold deposition (SEM images top row). The tissue-end of the wires is sharpened to limit tissue damage (SEM images bottom row), and electrochemically modified to optimise recording capabilities in the extracellular electrolyte (not shown, see Wray, 2017). (D) Splaying of an electrode bundle achieved by embedding micron size glass beads between the microwires. (E) Regular electrode spacing achieved by a sacrificial layer deposited onto the microwires before bundling (for details, see Hanna, 2019).

The thermal drawing process used for producing the wires allows great flexibility in the dimensions, and the inner / outer diameter ratio. Wire geometry can be adjusted by varying parameters, such as heating and pulling speed, and dimensions as small as 10 μm can be achieved, with nanoscale conductive cores (loisher et al., 2011; Yaman et al., 2011). The outer glass surface of the electrodes is highly consistent and smooth. The microscale composite wires are mechanically flexible, allowing wrapping as an approach to bundle electrodes, ensuring scalability (Badinter et al., 2010) in the order of hundreds of thousands to millions (Figure 2B).

Recording arrays from the microwires were prepared from spools to make bundles of 100-1000 wires. First, we embedded the wire bundle into PMMA polymer. Both the chip-end and the tissue-end of the bundle were finely polished with a grinder-polisher. Larger metal surfaces provide better contact interfaces, but wire diameter is a critical factor in tissue damage. To uncouple these two factors, and achieve optimal connection to the CMOS array, after polishing, we increased the exposed metal cores of the microwires by electrochemical metal deposition (Figure 2C). To ensure that local tissue displacement stays limited across the electrode array, the bundles were spaced by splaying (Figure 2D, Wray, 2017) or by incorporating a sacrificial spacer material during bundle fabrication (Figure 2E, Hanna, 2019).

The tissue-end was angle polished at 30° to achieve sharp wire tips, and the embedding material was removed by oxygen plasma, to free the last 2 mm of the tissue-end of the microwires from the embedding material (see *Materials and Methods* for details and alternatives).

### Electronics

While wires can be easily made in kilometres length and both electrochemical modification and polishing is inherently scalable and highly reproducible due to the standardized surface, using individual conventional amplifiers and operating amplifiers in general is not. CMOS integrated circuits on the other hand provide extensive scalability.

CMOS based high-density MEA chips are specifically designed for recording extracellular signals from brain tissue (Figure 3A). In addition to the high signal-to-noise ratio, further advantages are the capability for electrical stimulation and subselection of active pixels (Jones et al., 2011; Ferrea et al., 2012; Müller et al., 2015). Even higher channel count CMOS amplifier arrays are available in commercial IR sensors tailored to amplify small signals at high frame rate. In fact, they were shown to be in principle useable for slice MEA recordings (Scribner et al., 2007). We thus characterized an IR sensor, the Xenics Cheetah 640CL for its suitability to be used as a scalable amplifier platform for in vivo neural recordings. This IR camera is capable of frame rates of up to 200kHz (1.7 kHz full frame), which allows sufficiently high sampling rate to resolve neuronal action potentials. The CMOS readout circuit contains 327,680 capacitive transimpedance amplifiers with small feedback capacitances (7 fF, Neys et al., 2008). In the IR camera, the readout circuit is connected to photodiode layer using Indium bump bonding (Vermeiren et al., 2009). Individual amplifiers can thus be accessed through the Indium bumps (Figure 3B).

### Connecting bundle and chip

Contacting the microwire bundle to the CMOS chip in a scalable manner was achieved by a direct contacting approach (Figure 3C). Exploiting the high pixel count and density of the CMOS recording arrays, high-yield contacts were established by spacing the wires sufficiently (>100μm, see Hanna, 2019), and concurrently increasing the contacting area of individual wires (Figure 3D). With this approach, each wire is connected to a pixel with high probability, while single CMOS pixels are never connected to two wires. The chip end of the bundle was embedded into PMMA, and finely polished. To provide a sufficiently large contact surface for each electrode we electrochemically deposited 10 μm gold bumps onto the chip-end of the wires. The bundle was pressed against the ROIC (read-out circuit) using a custom designed press (Figure 3E). After removal of the bundle from the chip, the gold bumps showed deformation and indentations made by the CMOS surface, suggesting strong and stable contact. Connected pixels are sparsely distributed over the chip area (Figure 3F).

**Figure 3.**
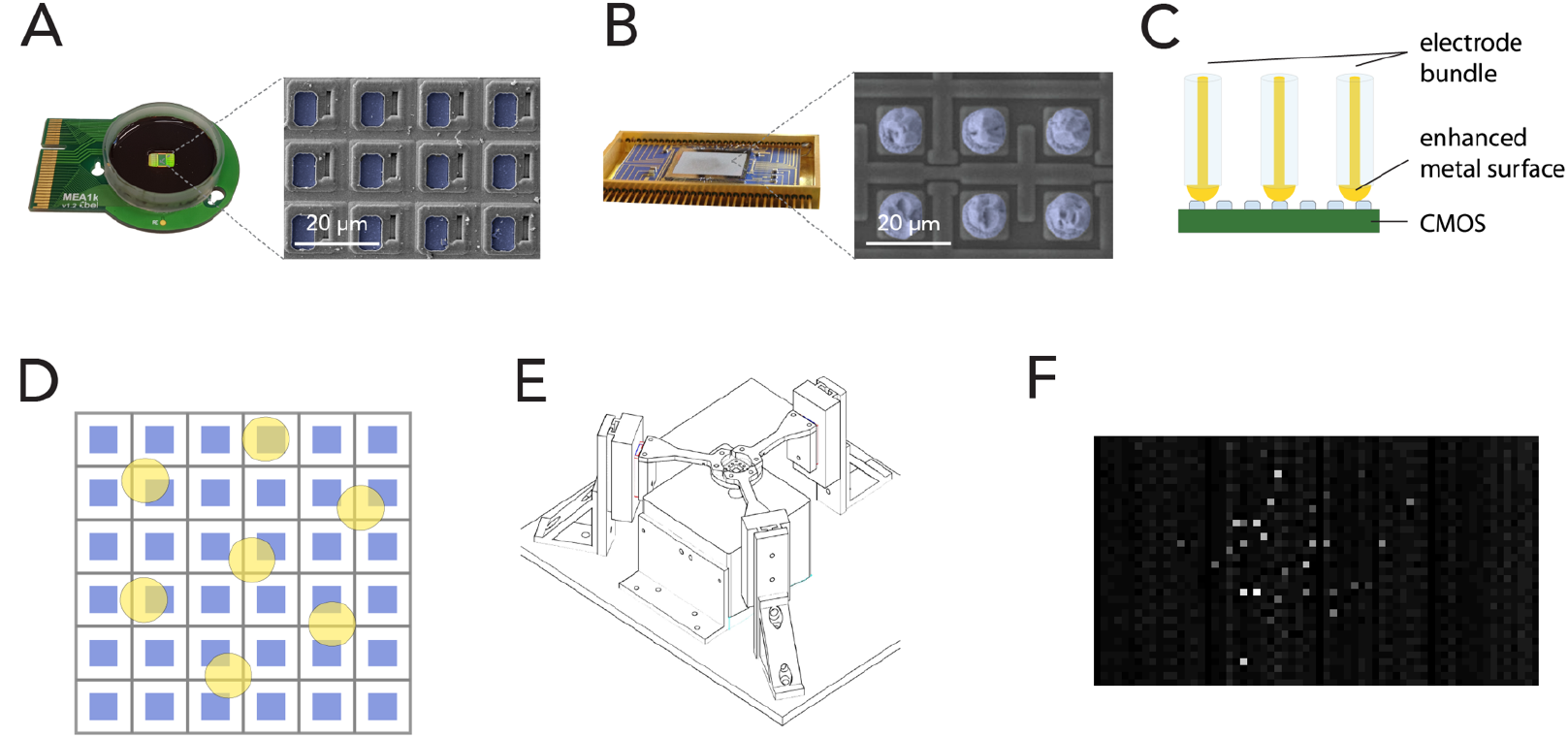
Scalable direct bonding of electrodes to CMOS pixels. (A) Mea1k, a high density, high signal-to-noise, switch-matrix based MEA chip (Müller et al., 2015). Inset: SEM of the chip surface. Blue pseudo-coloured areas denote the active Pt recording sites. (B) Commercial IR sensors provide another class of high pixel-count CMOS amplifier array, capable of sampling from >10^5^ pixels at rates of several kHz (here: Xenics Cheetah 640CL. Neys et al., 2008). Blue pseudocolour in the SEM image indicates Indium bumps on the amplifier input sites. (C) Direct contacting of electrode bundles to the conductive pixels is a highly scalable strategy to electrically interface bundles to electrode arrays. Increasing the size of the contacting metal surface of the microwires can be achieved in multiple ways and can enhance contact quality (e.g. Figure 2C). (D) High yield of contacts can be achieved without the alignment of electrode cores (yellow) and pixel recording areas (blue), by using sufficient spacing and large diameter electrode cores. (E) Alignment of the recording bundle and the flat CMOS array was achieved by a three-arm force-sensitive micromanipulator system. (F) A wire bundle of 200 electrodes connected to the amplifier array of an IR CMOS sensor. White pixels indicate pixels connected to electrodes.

### Signal-to-noise

Electrical signals in the extracellular space originating from single cells have typically amplitudes of few hundred microvolts in the close vicinity of the cell soma, and rapidly decay with distance. The signal amplitude picked up by any electrophysiological recording device is limited by the interfacial impedance between the solid-state conductor and the extracellular electrolyte (Figure 4A), and the properties of the amplifier. Signal loss and baseline noise levels in deep brain recordings are in most cases limited by stray capacitance. The stray capacitance depends on the insulating material and the ratio of the outer diameter (OD) and inner diameter (ID). Thus, glass/metal microwires with ODs of 10-20 μm and an ID of the metal core as small as 1 μm are theoretically predicted to have minimal stray capacitance due to the large OD/ID ratio provided pulled glass shares its electrical properties with bulk glass. To assess this, we immersed distinct lengths of wire into electrolyte, and measured the stray capacitance directly. The measurements confirmed a very low stray capacitance for the glass/metal composite wires (less than 80fF/mm, Figure 4B). While the small core and resulting large OD/ID ratio ensures minimal stray capacitance, the small exposed metal electrode surface at the tip of the electrode in principle could entail a large electrode impedance. To address this issue, we designed an electrodeposition protocol that lowers the specific impedance of the electrode-electrolyte interface and can be performed on an arbitrary number of electrodes simultaneously (see Wray, 2017).

To probe the ability of CMOS pixel amplifiers to record small voltage fluctuations expected during recordings, we connected bundles, electrochemically modified as described above, to two types of CMOS chips (MEA1k: Müller et al., 2015, and Xenics Cheetah 640-CL), and submerged the tissue-end into a voltage-clamped bath. In order to compensate for the reference voltage offset of the chips (1.65V for the MEA1k and 2.3V for the Cheetah 640-CL, Neys et al., 2008), the bath was voltage-clamped using custom electronics to the appropriate offset (see Methods). For the Cheetah CMOS system, pixels were activated by large (2-3 V) voltage steps to break any potential residual oxide layer (Gemme et al., 2001). We characterized the electronics by measuring the response to 1 mV square and sinusoidal voltage signals applied to a bath of saline, and found that individual pixels are capable of recording voltage signals with a noise floor of 17.4 ± 1.2 μV (RMS) on the MEA1k chip, and 58.2 ± 21.5 μV for the Cheetah 640-CL. This resulted in signal-to-noise ratios of up to 29 and 9 respectively, given 500 μV spike amplitudes, which were recorded reliably in the mouse brain. After subtracting correlated noise, the residual RMS noise was 6.5 ± 2.6 μV in connected MEA1k pixels, suggesting that noise levels could be further reduced by shielding.

**Figure 4.**
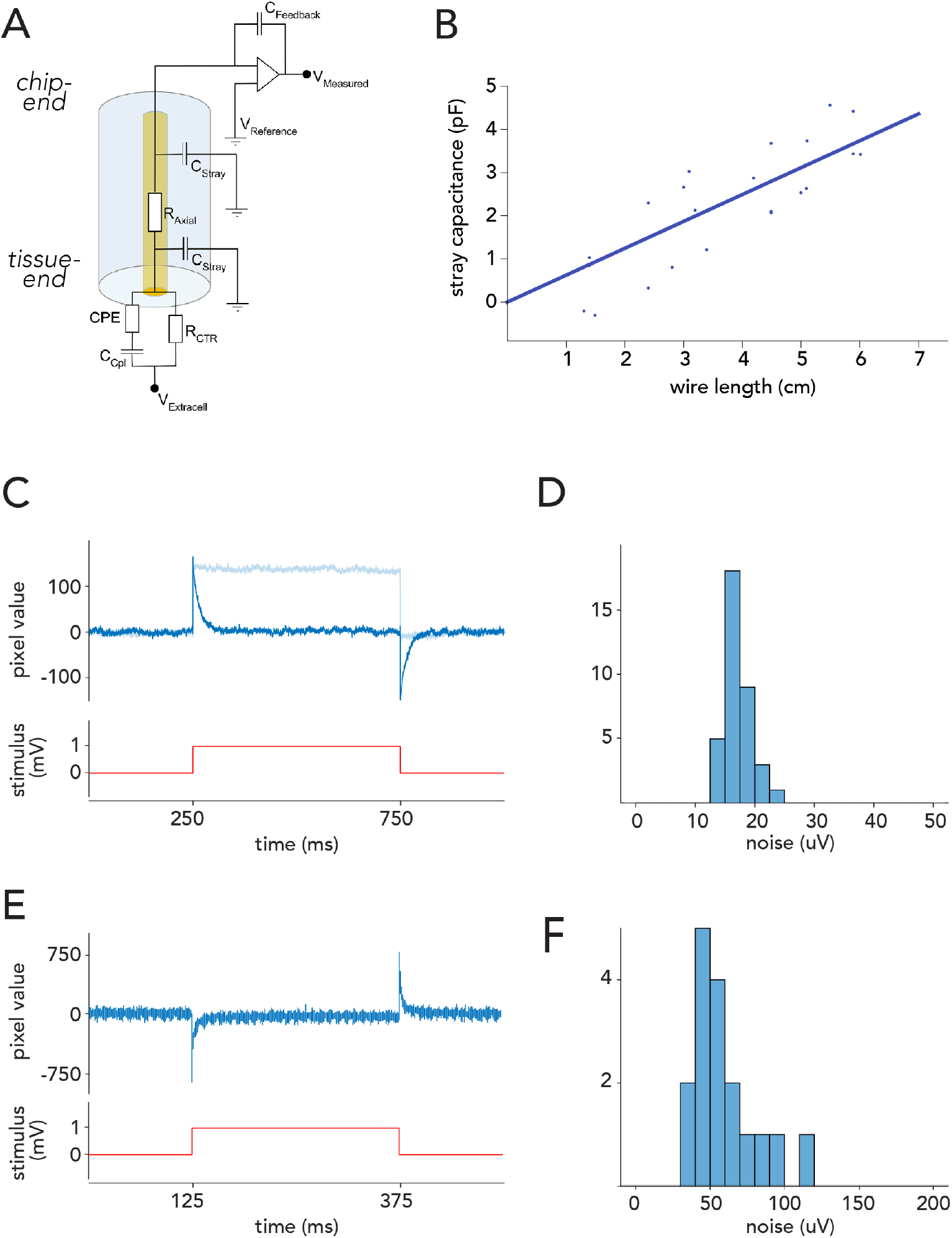
Noise levels in voltage recordings with CHIME. (A) Electrical model of the recording configuration (depicted for a capacitive transimpedance amplifier derived from IR sensors (Vermeiren et al., 2009)). The axial resistance of the metal core microwires (R_Axial_) is low (<300 Ω/cm). Signal loss is mainly determined by the stray capacitance (C_Stray_). Signal amplitudes depend on the feedback capacitance of the amplifier (C_Feedback_), and the degree of interfacial impedance, primarily the coupling capacitance (C_Cpl_). (B) Stray capacitance for a glass/gold wire with outer diameter of 28.3 μm and inner diameter of 1.1 μm. Stray capacitance scales linearly with wire length and is low (<1pF/cm) due to the high outer to inner diameter ratio. (C) Response of one pixel of a Mea1k switch-matrix based MEA (Müller et al., 2015) to a 1 mV voltage step applied to the bath, recorded through a connected electrode. On-chip high-pass filter: 300 Hz (dark blue), 1 Hz (light blue) (D) RMS noise level distribution in connected Mea1k pixels. (E) Voltage response of a Cheetah 640CL IR camera ROIC pixel (Neys et al., 2008). Due to the inverting amplifier circuit, responses are reversed. (F) Noise level distribution in connected IR camera CMOS pixels.

### Histology

The fundamental limit to scalability of electrodes is often considered to be tissue damage Marblestone et al., 2013). The size of the implant is strongly correlated with the extent of damage, and it has been suggested that small wires <20 μm would not cause significant damage to the brain tissue (Kozai et al., 2012; Guitchounts et al., 2013). Extensive surface tension due to penetration force, increased pressure due to volume displacement and potential direct effect of friction of the moving electrode are thought to be the underlying reasons for tissue injury (Edell et al., 1992; Subbaroyan et al., 2005). Beyond the direct damage to neurons, the rupture of vessels is a strong indicator of tissue damage, and a predictor of subsequent degradation of recordings (Kozai et al., 2012; Prodanov and Delbeke, 2016). Histology after insertion experiments with single CHIME electrodes indicated minimal damage to the vasculature (see Wray, 2017). In case of large channel-count electrode bundles, a high density of recording sites is preferred. For bundle implantations, there is likely a trade-off between the density of recording sites and amount of damage caused, making the spacing between shanks an important factor. Considering these factors, we tested the tissue compatibility of bundles of 500 wires with an average inter-wire spacing of around 100 μm.

**Figure 5.**
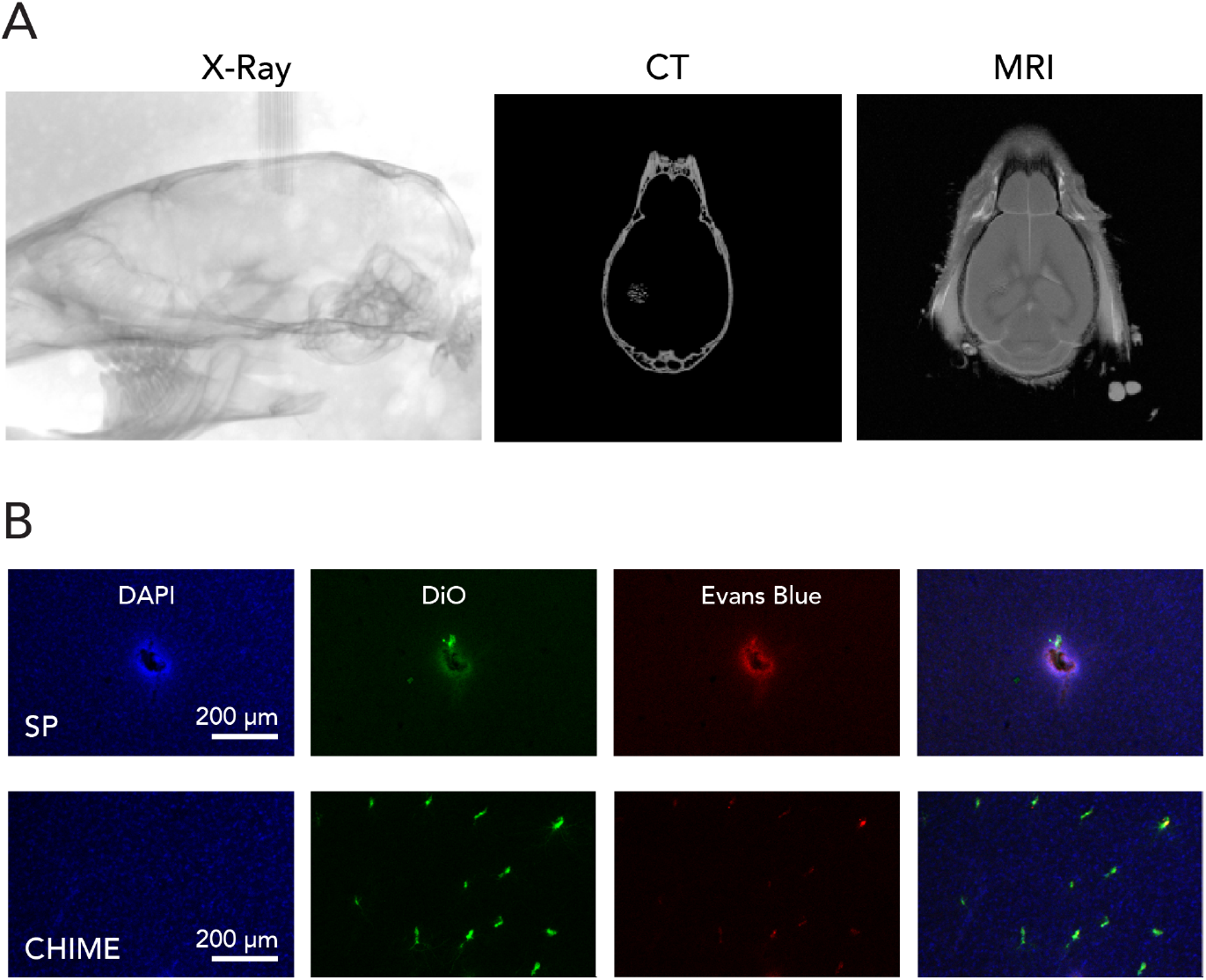
Brain tissue compatibility of CHIME microwire bundles. (A) X-ray microscopy images of a CHIME bundle inserted into mouse cerebrum, and correlated MRI image. (B) Comparative histology of the acute blood-brain-barrier (BBB) injury upon standard silicone probe insertion (SP) and insertion of a CHIME electrode bundle. Evans Blue was injected into the bloodstream, which is only detectable upon BBB damage (red). For the tracking of probe locations, electrodes were dipped into DiO solution before insertion (green).

The tissue-end of the wires were sharpened and prepared as described above and inserted into the cortical tissue after the removal of the dura mater. Wires inserted readily into the tissue, without buckling, as expected from estimation of the buckling force for the glass-metal microwires (≈300 μN for a mm long microwire). For the localization of recording sites, the electrode bundle was visualized inside the skull (Figure 5A) by a 3D X-ray. Individual microwires could be visualized. Apart from stereotaxic coordinates relative to bone landmarks, further delineation of the anatomical location of each recording site is possible with correlated MRI (Figure 5A).

We assessed the extent of damage to the blood-brain-barrier potentially caused by spaced dense CHIME bundles, by injecting Evans Blue into the bloodstream, followed by insertion of the electrodes into the brain, and perfusion fixation. As Evans Blue is bound to albumin, its presence in the brain tissue would indicate damage to the blood-brain-barrier (Rawson, 1943; Belayev et al., 1996). The results confirmed minimal damage upon insertion of sharp 20 μm glass/metal fibres with an average spacing of 100 μm, markedly less compared to the extravasation caused by a standard silicone probe (Figure 5B).

### Neuronal recordings with CHIME

Finally, we performed *in-vivo* neuronal recordings using the here described CHIME assembly with both chips (Figure 6). The switch-matrix based CMOS MEA architecture (Müller et al., 2015) has the advantage that recorded channels can be pre-selected (1024 channels from 26,400), thus it is not necessary to stream and record pixels not connected to recording electrodes. Additionally, programmable high-pass filter, DC-offset cancellation and voltage or current stimulation through individual pixels/electrodes is available. In comparison, a ROIC repurposed from a CMOS IR camera (Xenics Cheetah 640-CL) provides 327,680 simultaneously recorded pixel amplifiers. As offset compensation or filtering is not possible in this architecture, large drifts more often lead to pixel saturation. In case of the IR CMOS camera, voltage traces can be extracted from connected pixels post-hoc from the video stream recorded by a DVR system.

We recorded neuronal activity from the mouse olfactory bulb with CHIME on both CMOS systems. With the MEA1k, large-amplitude LFP signals were found on all inserted electrodes (Figure 6A), and high signal-to-noise single unit spikes (Figure 6B) were recorded. The IR-camera chip provided lower amplitude LFP signals, due to the high-pass filtering characteristic of the pixel amplifier (Figure 6C), but well isolated single unit spikes could be recorded on individual channels (Figure 6D). The CHIME technique provides a means for scalable, high-temporal resolution, parallel recordings from mammalian brain circuits with high lateral density (Figure 6E).

**Figure 6.**
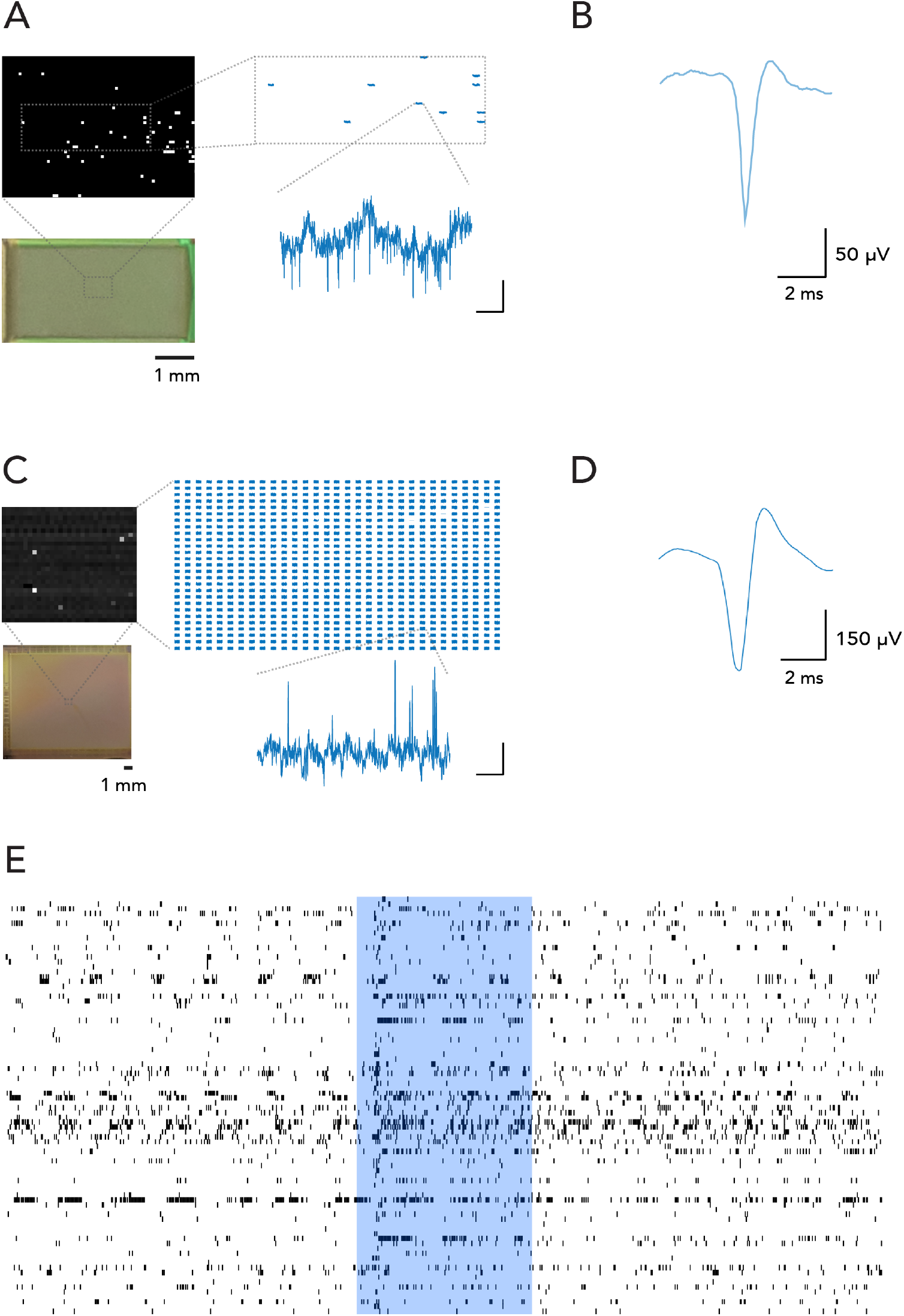
Large-scale, parallel electrophysiological recordings with CHIME. (A) Switch-matrix based MEA chips allow recording selectively from the pixels connected to electrodes. The inset shows recorded traces from the frame. Only pixels connected to electrodes are pre-selected and recorded. Voltage trace scale bars: 5 ms and 200 μV. (B) Average spike waveform recorded from the mouse main olfactory bulb with the switch-matrix MEA chip (Müller et al., 2015). (C) CMOS IR-camera chips (Neys et al., 2008) provide a frame stream from a large number of pixels. All pixels are recorded in a pre-selected rectangular region, of which only a subset is connected to electrodes. Scale bars: 5 ms and 200 μV. (D) Average spike waveform recorded from the mouse main olfactory bulb with the IR camera chip. (E) Spike events recorded from 86 channels in the mouse main olfactory bulb. The blue area indicates the timing of a 1 s long odour stimulus.

## Discussion

Here we have used flexible, glass insulated, metal microwires connected to a CMOS readout circuit to perform electrical recordings from neurons. The described technique, CMOS Hosted In-vivo MicroElectrode (CHIME) recording, promises to drastically increase the number of simultaneously recordable neurons. Recording sites can be arranged in arbitrary locations, allowing the experimenter to query neuronal networks *in-vivo*, such that the arrangement is driven by the addressed functional question rather than the physical limitations of the recording system.

What are the limits to the scalability? At the level of the amplifier array, the number of channels in CMOS chips increases exponentially (Moore, 2006), and the latest designs already substantially surpassed the ≈300.000 channels available in the Xenics Cheetah 640CL camera. Imaging chips have the potential to continue to scale following Moore’s law, reaching 10^7^-10^8^ channels in the near future. MEA chips, which are routinely used to record from cultured cells, brain slices or flat preparations such as the retina, also show rapid development, and chips with significantly larger pixel counts are anticipated in the following years. While the connectivity between electrodes and chip did pose development challenges, we have not seen any decrease in the efficiency of connection in larger bundles (Hanna, 2019). At the scale of the electrode bundles, achieving a sufficiently flat surface for reliable connection is not too demanding, and far from the standards needed for e.g. silicon wafers. Due to the flexibility of the wires, the fabrication of electrode bundles is inherently scalable. A potential limit to channel counts could result from tissue damage. We have shown that wires at the 15-20 μm scale cause minimal damage to the tissue, supporting the notion that damage to the tissue depends non-linearly on the size of the electrode. We also have seen that blood vessels avoid the electrode tip during penetration (Wray, 2017), which prevents bleeding and subsequent degradation. While we have investigated acute damage, the small size of wires and smooth surface also provides mechanical compliance (Seymour and Kipke, 2007; Etemadi et al., 2016; Luan et al., 2017) and thus likely enhanced biocompatibility. With increasing number of wires, we expect at some point tissue displacement to become the limiting factor. As a solution, wires could be made smaller. With smaller wire diameters, however, at some point the tip impedance becomes too high, impeding signal strength. As a solution, novel materials could be used on the electrode-electrolyte interface which further increase the specific surface area and improve coupling (Kim et al., 2010; Prasad and Sanchez, 2012; Baneyx and Park, 2013; Chung et al., 2015). Considering the input impedance of the amplifiers, when using 15-20 μm wires, tip impedance is magnitudes lower than what would become a limiting factor, and further improvements can be made for smaller probes.

With small wires, another challenge is posed by buckling. The critical buckling force decreases with the fourth power of the radius, therefore with decreasing size, the force that the wire can apply onto the tip drops significantly. This causes problems if the maximal force is below the force needed for insertion. Measurements have been made for different types of electrodes (Sharp et al., 2009; Casanova et al., 2014; Fekete et al., 2015), and recently a quantitative study was undertaken to understand the penetration mechanics of microwires specifically (Obaid et al., 2018). In our experiments, 15-20 μm wires could be inserted into the cortex several millimetres deep without support. For smaller wires, a possible solution is the use of temporary coating agents (Foley et al., 2009; Lewitus et al., 2011; Etemadi et al., 2016; Singh et al., 2016). Wire fabrication with the Taylor-Ulitovsky method allows direct fabrication of approx. 10 μm wires. Smaller dimensions can be achieved by using a three-component pre-form, with a conductive core, glass inner cladding and etchable glass outer cladding. The etchable glass could then be specifically removed by acids during bundle fabrication analogously to the fabrication of nanochannel array glass (Tonucci et al., 1992) or leached fibre bundles (Gerstner et al., 2004). Another approach is to use thermal drawing to produce tapering in the electrodes (Ioisher et al., 2011; Yaman et al., 2011).

Flexible wires do not necessarily follow a perfectly straight trajectory through the tissue; while this allows them to e.g. avoid blood vessels, it also renders positioning more challenging compared to larger or rigid probes. Approximate electrode locations can be adjusted by the spacing of wires during bundle fabrication, however the recording sites are not necessarily homogeneously distributed. Nevertheless, when the number and average density of recordings sites allows the recording of virtually all units through multiple channels, this arrangement has less importance. Direct localization of the wires can be achieved by high-resolution X-ray tomography (Figure 5A), combined with other methods to identify specific brain areas and layers. Another possible approach to localize recording sites is the use of “electrical imaging”, whereby the analysis of noise and signal correlations between channels can be exploited to predict their relative location (Cybulski et al., 2015).

While glass wires provide many advantages including high insulation, smoothness, other types of wires could be used as they get optimized and become available in large quantities. Carbon fibres (Guitchounts et al., 2013) have the advantage of stiffness, and good recording properties, while fabrication processes are as of now less scalable, and have a higher stray capacitance and less flexibility. Plastic wires can also be used (Canales et al., 2015) and have the advantage of great flexibility, and tissue compatibility. On the other hand, the metals, which can be used at the low temperatures currently needed for plastic fibre drawing (e.g. tin), provide a less advantageous substrate for electrochemical modifications compared to higher melting-point metals, such as gold or copper, and might require larger core diameters (and thus increased stray loss) due to reduced electrical conductivity.

As with other large-scale approaches, data handling and, in particular, data analysis will be a key challenge. Identifying and isolating spiking activity from individual neurons is an important requirement and is typically resolved by clustering based on waveform. Large amplitude spikes are both more informative on the waveform and originate from a smaller volume around the electrode. Therefore, the extent of local damage around the electrode has a strong impact on the reliability of the assignment of spikes to individual neurons, which points to the advantage of CHIME, which show minimal local damage.

The system described here is currently designed for head-fixed experiments. However, the extremely low stray capacitance (< 100 fF / mm) makes it possible to record through long (10s of cm) wire bundles, thus immediately allowing tethering. In addition, the CMOS chip itself is relatively small, it is thus furthermore conceivable – at least in larger mammals – to abandon tethering altogether, move the active electronics to a location outside the skull and perform signal processing, compression and ultimately wireless signal transmission there.

In summary, combining microwire bundles and integrated circuit arrays allows for a highly scalable neuronal recording approach, essentially tying the progress of electrical neuronal recording to the rapid scaling of silicon microfabrication. Moreover, our approach of employing bundles of minimally invasive, highly insulated and functionalized microwires as a means to extend a 2-dimensional CMOS architecture into the 3^rd^ dimension can be translated to other integrated circuit arrays such as electrical stimulation devices and thus might provide a pathway for high bandwidth bidirectional interfacing with neural circuitry.

## Materials and Methods

Glass ensheathed gold microwires (ELIRI, S.A., Moldova) were obtained as continuous threads of various lengths (50-500 m) on individual spools. Bundles were created by wrapping microwire from spools with a modified winding machine (Optima 1100, Synthesis, India). The bundle electrodes were then sealed by applying 2-component epoxy resin (Uhu, Germany) on both sides of the bundle. CB 509 polymer (Agar Scientific, UK) was heated to 126°C in a metal crucible within a hot plate desiccator and then pumped to 0.9 bar for 10 minutes to remove bubbles from the molten CB. The chip end of each electrode was dipped in molten CB 20 times, removed from the crucible, and hung up for a few minutes to harden. The bundles were centered within a polypropylene tube (2 mL syringe) and the chip end was back filled with molten CB by pouring from a ladle to fuse it within the tube. The chip end was polished flat by fixing the supporting tube in a custom polishing holder. This holder was designed to fit the pneumatically powered head of a rotating polishing machine (Metaserv 250 with Vector Power Head, Buehler, IL, USA) and could support 4 electrodes simultaneously. The electrodes were first balanced manually, semi-automatically polished at grit 240 (ANSI), then re-aligned and polished on grit 320 and 600 slurries. The electrode face was then washed with distilled water and inspected with a dissection microscope. The tissue end was embedded in dental cement (Paladur, Kuzler, Germany) by filling the end of another polypropy-lene tube supported in a mold with freshly mixed liquid. The cement was allowed to harden for 10 minutes. The cement embedded tissue end was flat polished as described previously.

The embedded polished wires at the tissue end were electrically connected by sputtering gold for 170 seconds with a sputterer (Agar Scientific, UK). Next, an insulated macroscopic wire was connected to the sputtered surface with conductive silver epoxy (MG Chemicals, Canada) which was allowed to harden overnight. The silver epoxy was then coated with epoxy resin to insulate and mechanically support the macroscopic wire. Gold bumps were electrodeposited on the chip end to increase the contact surface area of each wire. ~1 cm of the tissue end was cut off with a hot scalpel blade. The tissue end was embedded in CB but was supported in a polypropylene tube cut at a 30° angle. The tissue end was polished similar to the chip end, but with a custom polishing holder which held electrodes at a 30° angle to result in sharp “needle” tips for simplified tissue insertion. The tissue end was freed from its CB embedding by removing the CB with a solvent stripper (509-S, Agar Scientific, UK). Removal of CB residue was achieved after immersing in 150 mL of stirred stripper for at least 16 hours with the stripper replaced at 3 different intervals. The freed end was then washed in distilled water. An alternative approach for increasing the metal surface on the chip-end by glass etching is discussed by Hanna (2019).

An alternative method was used in a subset of experiments, where both ends of the bundle were embedded into dental cement. After polishing, the tissue end was freed by oxygen plasma ashing (5 hours at 100W, SPI Plasma Prep III).

Gold and iridium oxide (IrOx) was electrodeposited on the tissue end as described in Wray, 2017. To electrically connect to the chip end, the embedded gold bumps were pressed into a tightly fitting connector filled with a paste of carbon powder (Sigma) and mineral oil. This created a temporary electrical connection with contact resistance of 100-500 Ω per wire. After electrochemical deposition, the chip end was freed from its CB embedding and the carbon paste was washed off with CB stripper.

The double functionalized electrode bundle freed from embedding was then mounted to a flat length of polystyrene (end of cuvette stirrer) with 2 part epoxy. ~2-5 mm of the tissue end was left freed on both sides of the bundle electrode. The polystyrene shank was then mounted with 2 part epoxy to a custom made holder fit for the chip alignment setup. Recordings were performed with either a MEA1k switch-matrix MEA chip (Müller et al., 2015) or a modified Cheetah 640 CL infrared camera (Xenics, Leuven, Belgium). The photosensitive layer was omitted by the manufacturer, so that individual amplifiers at each pixel of the camera chip could be directly accessed via indium bumps. The ~10 μm pixels have a pitch of 20 μm in a 640×512 array. The Cheetah 640 CL can operate in high and low gain mode; in the high gain mode used during recordings the feedback capacitance is 6.73 fF, sensitivity is 22.8 μV/electron, and the saturating charge is ~75 000 electrons with 192 electrons of noise (Neys et al., 2008). The reference voltage for the camera (virtual ground of the transimpedance amplifier of every pixel) was approximately 2.3 V. Throughout this work, the camera was operated in integrate-while-read mode where each frame is integrated while the previous frame is readout through the camera link ports. Depending on the selected window of pixels, recordings took place at 1.7 kHz (full frame) to 200 kHz (smallest frame). A DVR System (CORE, IO Industries, Canada) was used to continuously record data at up to 2 GB/s.

For the characterisation of the pixel amplifier, an optically isolated, low noise voltage source was custom developed (Wray, 2017) to move the pixel within saturation limits by raising the voltage ~1.65/2.3 V above ground and then applying voltage waveforms with < 50 μV noise into a bath containing 0.9% NaCl solution through a standard Ag/Cl electrode.

Resistance and stray capacitance of glass ensheathed microwires were measured with an LCR meter (LCR-821, GW-Instek, Taiwan). A 3D printed chamber supported a microwire connected to 2 ports filled with conductive liquid Galinstan. The resistance across the 7.5 cm wire was measured with both ends of the wire connected to a Galinstan port. An external electrode (copper wire) was immersed along with the connected microwire in PBS solution for various lengths and the capacitance was measured from the external electrode (in grounded solution surrounding the microwire) to one of the galinstan ports.

The chip-end of the electrode bundles can b aligned and pressed onto the chip using different solutions: a custom 3-arm press system built on 3 micromanipulators (Luigs & Neumann, Germany) joined with a magnetically attached flexible centrepiece and independent strain measurements on each arm (Wray, 2017) was used for recordings with the Cheetah camera chip, and a screw-based passive alignment system (Hanna, 2019) with the Mea1K. Pixels contacted to the chip were detected by significant deviations in the average pixel value (mean ± 2 SD).

C57BL/6 mice aged 4-6 weeks were used in anesthetized recordings. To expose the olfactory bulb (OB), craniotomy surgeries were performed as described before (Kollo et al., 2014). For surgery, mice were anesthetized using ketamine (100 mg per kg of body weight) and xylazine (20 mg per kg for induction and 10 mg per kg for maintenance) administered intraperitoneally and supplemented as required. Body temperature was kept at 37°C with a feedback regulated heating pad (FHC, ME, USA). A custom-made steel headplate was fixed to the parietal and interparietal bone plates with dental cement. A 2×3 mm craniotomy was drilled over olfactory bulb. The chamber surrounding the craniotomy was filled with Ringer’s solution to prevent it from drying out. Next, a Agl/AgCl reference electrode was placed in the meniscus of Ringer’s solution.

After surgery the anesthetized mouse was brought directly to the recording setup and kept under anaesthesia with intraperitoneal supplementation of ketamine/xylazine as needed. The mouse was placed on a platform mounted on a three-axis manipulator below the recording device with the electrode bundle facing downwards. Respiration was monitored by a piezoelectric sensor and body temperature maintained at 37°C. Using a small web camera (Raspberry Pi, Raspberry Pi Foundation, UK) and mirror, the bundle electrode was aligned over the craniotomy. The reference electrode in the meniscus over the brain was lifted to the reference voltage of the camera using a low-noise voltage source (see above). Wire connectivity and noise were roughly assessed in the meniscus before insertion.

All animal experiments were approved by the local ethics panel and the UK Home Office under the Animals (Scientific Procedures) Act 1986.

### Histology

After the craniotomy was performed, 0.2ml of 0.5% Evans Blue was injected into the tail vein. Insertion of the bundle electrode occurred within 30 minutes of the tail vein injection. Prior to bundle insertion, wires were dipped into SP-DiO (Molecular Probes, OR, USA) and allowed to dry. The bundle electrode was inserted into the cortex. Immediately after the wire bundle was removed the mouse was perfused with ice cold 4% PFA, the brain was harvested and stored in 4% PFA overnight. Using a Vibratome (Leica, Germany), the brain was sliced into 100 μm horizontal sections. These sections were stored in 1x PBS during slicing. The slices were stained with DAPI using a 1:1000 DAPI:PBS wash for 10 minutes and were then transferred to fresh PBS. Slices were mounted and coverslips were sealed. Imaging was completed on a confocal microscope (Leica SP5).

### CT/MRI

For CT/MRI images, after performing a craniotomy as described above, a bundle was inserted into the brain with a micromanipulator (SM7, Luigs and Neumann). The mouse was then given an overdose of the anaesthetic, subsequently decapitated and the whole head was embedded into a block of epoxy resin (Uhu, Germany) and attached to the microscope specimen holder. The whole head was imaged with a microCT (XRadia 510 Versa, Zeiss, Germany) at 30kV (50.5 um pixels, 40-minute scan) and then the region of interest at the tissue end of the inserted bundle in higher resolution (3.98 um pixels, 10-hour scan). Imaris software was used to extract and semi-manually delineate the features (bundle, skull and jaw bone, and the soft tissue).

## Acknowledgements

We thank Martyn Stopps for help with electronic design, Isabell Whiteley for technical help with histological processing, Jan Vermeiren for advice on the Cheetah camera CMOS, Carles Bosch for help with CT scans, Anatolii Ioisher for advice on glass-metal microwires, and Konrad Kording for intense discussion throughout the project. We also thank Lucy Collison and the EM STP for help with SEM imaging, Bernard Siow for help in MRI and CT imaging, the Making Lab and the BRF for technical help. This work was supported by the Francis Crick Institute which receives its core funding from Cancer Research UK (FC001153), the UK Medical Research Council (FC001153), and the Wellcome Trust (FC001153), an HFSP grant to A.T.S. and N.A.M. (RGP 00048/2013), an NIH BRAIN initiative grant to A.T.S. and N.A.M. and Konrad Kording (1U01NS094248-01), and the Medical Research Council (MC_UP_1202/5). Andreas Schaefer is a Wellcome Trust investigator (110174/Z/15/Z).

